# Electroencephalography-derived functional connectivity in sensorimotor networks in Stroke and Multiple Sclerosis Fatigue

**DOI:** 10.1101/2022.03.16.484592

**Authors:** Chi-Hsu Wu, William De Doncker, Pierpaolo Croce, Massimo Bertoli, Franca Tecchio, Annapoorna Kuppuswamy

## Abstract

A common mechanism of altered sensory processing is the basis of chronic fatigue in neurological disorders. Here we test the hypothesis ‘Altered connectivity in sensory networks underlies chronic fatigue in stroke and multiple sclerosis’.

In 46 non-depressed, minimally impaired stroke survivors (n=29) and multiple sclerosis patients (n=17), median disease duration of 5 years, resting state neuronal activity was measured using 64-channel electroencephalography. Graph theory-based network analysis measure of functional connectivity (small-world index) was calculated in right and left motor (Brodmann areas 4, 6, 8, 9, 24 and 32) and somatosensory (Brodmann areas 1, 2, 3, 5, 7, 40 and 43) networks, in 5 frequency bands: delta, theta, alpha, beta and gamma. Fatigue was measured using Fatigue Severity Scale (Stroke) and modified Fatigue Impact Scale (MS), with scores of >4 (FSS) and >38 (mFIS), defined as high fatigue.

Both stroke survivors and multiple sclerosis patients with high fatigue showed significantly more small-worldness in the right sensory networks in the beta band frequency. Additionally, only in stroke survivors with high fatigue, there was decreased small-worldness in the left motor network in the delta and theta bands.

Altered sensory network connectivity is common to both stroke and MS fatigue, indicating impaired sensory processing as a disease-independent mechanism of chronic fatigue in neurological conditions. Furthermore, such difference in functional connectivity emerges in beta band activity, further strengthening the idea of altered sensorimotor processing as the basis of chronic neurological fatigue.

## Introduction

Stroke, a result of vascular insufficiency to neurons; and multiple sclerosis (MS), a neurodegenerative disease, both present with fatigue as a significant symptom. In both conditions, the disease processes do not explain the reported levels of fatigue^1,2^. We have previously proposed a disease-independent mechanism of fatigue in neurological disorders that revolves around the idea of poor suppression of anticipated sensory information^3,4^. During movement, poor suppression of muscle sensory afferents results in assigning high effort to simple tasks^5^. During visual and auditory perception, lack of distractor suppression leads to fatigue. From the view of predictive brain processing, such task related high effort, informs sensory priors at rest, explaining how one can experience fatigue at rest without performing any activity.

From a neuronal network functioning point of view, behavioral development mirrors structural and functional changes of the networks, which persist even in resting state^6^. In other words, neuronal networks at rest express features that keep trace of their ability to perform the required behaviour^7–11^. These structural and functional features of the networks at rest also display alterations related to chronic symptoms^12^. Consistently, both in post-stroke fatigue and MS fatigue, neurophysiological^13–18^ and behavioural^19–21^ findings support an altered resting state, where abnormal sensory network activity is a common driver of chronic fatigue across both neurological conditions.

Reduced corticospinal excitability at rest, in post-stroke fatigue is driven by diminished inputs to the motor cortex^13^, such as sensory input from muscles, which directly modulate corticospinal excitability at rest^22^. Likewise, immediately prior to movement, there is reduced modulation of corticospinal excitability both in stroke^15^ and MS ^17^ which can be explained by poor integration of sensory inputs with corticospinal output^23^. Reports of greater perceived effort related to muscle activation both in stroke^21^ and MS^24^ fatigue, suggests altered processing of sensory input from the muscles. Moreover, those with stroke report a ‘heaviness’ associated with their limbs which significantly correlates with self-reported fatigue but not muscle weakness^20^ suggesting that fatigue might be related to abnormal sensations related to resting muscle tone. In MS, resting state connectivity studies show altered functional connectivity in sensory networks^25^ with targeted neuromodulation of somatosensory networks significantly reducing fatigue^12,26^. Therefore, a common mechanism of pathological fatigue may be underpinned by sensory network activity alteration, which can be sensed even at rest.

Ensembles of neurons that fire at specific frequencies and communicate with each other by synchronising their firing, comprise a neuronal network. To understand a network’s activity, the strength of synchronicity between various nodes is mapped using functional connectivity methods^27^. Functional connectivity is defined as the temporal correlation or dependency between distinct neuronal groups and areas^28,29^. Such temporal correlation occurs in various frequency bands, with low frequencies associated with arousal, mid-range frequencies related to sensorimotor activity, and high frequencies representing higher order functions such as error detection and learning. With pathological fatigue in stroke and MS proposed to be a problem of sensorimotor control, specifically arising from processing of incoming muscle related sensory information, we anticipated a fatigue related modulation of beta band frequency.

Therefore, in this resting state study we tested the hypothesis ‘Fatigue in stroke and multiple sclerosis shares alterations of functional connectivity in somatosensory networks’.

## Methods

### Participants

This was a cross-sectional observational study approved by the London Bromley Research Ethics Committee (REC reference number: 16/LO/0714) and the Ethical Committee of ‘S. Giovanni Calibita’ Fatebenefratelli Hospital Rome (n.22/2012). Stroke survivors were recruited and tested at the Institute of Neurology, London, UK while MS patients were recruited and tested at the Fatebenefratelli Isola Tiberina Hospital.

Stroke survivors were recruited via the Clinical Research Network from the University College NHS Trust Hospital (UCLH), a departmental Stroke Database and from the community. All stroke survivors were screened prior to the study based on the following criteria: first-time ischaemic or haemorrhagic stroke; stroke occurred at least 3 months prior to the study; no clinical diagnosis of any other neurological disorder; physically well recovered following their stroke defined as grip strength and manual dexterity of the affected hand being at least 60% of the unaffected hand assessed using a hand-held dynamometer and the nine-hole peg test (NHPT) respectively; not taking anti-depressants or any other medication that has a direct effect on the central nervous system; not clinically depressed with depression scores ≤ 11 assessed using the Hospital Anxiety and Depression Scale (HADS). HADS is a 14-item questionnaire with a depression and anxiety subscale with a score of 0 to 7 for either subscale regarded as being in the normal range, while a score of 11 or higher indicating probable presence of the mood disorder^30^.

MS patients were recruited from the MS Centre of the Fatebenefratelli Isola Tiberina Neuroscience Department and were screened prior to the study based on the following criteria: absence of clinical relapse or radiological evidence of disease activity over the preceding three months; no clinical diagnosis of epilepsy or any other central/peripheral nervous system comorbidity; no coexistence of other condition(s) potentially associated with fatigue (i.e. anaemia, pregnancy); low clinical disability (Expanded Disability Status Scale, EDSS ≤ 2); not taking anti-depressants, psychoactive or medication that may affect the level of fatigue, depression and anxiety within the preceding three months; not clinically depressed with depression scores <15 assessed using the Beck Depression Inventory (BDI), a 21-item questionnaire with a score ranging from 0 to 63^31^.

Overall, twenty-nine stroke survivors and seventeen MS patients took part in the study (Table 1 and 2) and provided written informed consent in accordance with the Declaration of Helsinki.

### Fatigue

A measure of trait fatigue was captured at the start of the study, which represents the experience and impact of fatigue on day to day living for a pre-determined time leading up to the day of testing. In the stroke cohort, trait fatigue was quantified using the FSS-7, a seven-item questionnaire asking for ratings of fatigue ranging from one to seven (strongly disagree to strongly agree) over the preceding week from the day of administration. An average score of one indicates no fatigue while an average score of seven indicates very severe fatigue^32^. Stroke survivors were subsequently divided into two groups, high and low fatigue, based on their FSS-7 scores. A cut-off score of less than four on the FSS-7 was classified as low fatigue and a score equal or greater than four was classified as high fatigue^33^. In the MS cohort, trait fatigue was quantified using the modified fatigue impact scale (mFIS), a 21-item multidimensional questionnaire that measures the physical, cognitive and psychosocial impact of fatigue using a five-point ordinal scale^34^. A total score of zero indicates no fatigue while a total score of 84 indicates very severe fatigue. A cut-off score of 38 is commonly used to define high fatigue^35^. The MS patients recruited in this study were all classified as high fatigue.

### EEG recording

Whole-scalp electroencephalography (EEG) data was recorded using 64-channel systems, ActiCap, Herrsching, Germany, and a BrainAmp (stroke cohort) or actiChamp, BrainProducts, Gilching, Germany (MS cohort), while participants were at rest, with their eyes open and focusing on a fixation cross displayed on a monitor. The duration of the recording in the stroke cohort was seven minutes long while that of the MS cohort was three minutes. The 64 electrodes were positioned on the cap in accordance with the 10-20 international EEG electrode array. During online recordings, channels FCz and AFz were used as the reference and ground respectively. Impedances were kept below 10 kΩ throughout the recording. The EEG signal was sampled at 1 kHz and visualized online using the BrainVision Recorder Software (BrainVision Recorder, Version 1.21.0102 Brain Products GmbH, Gilching, Germany).

### EEG Analysis

EEG analyses were performed with a combination of EEGLAB^36^ and custom Matlab scripts. EEG data was down-sampled to 250 Hz and then band-pass filtered from 0.1 to 47 Hz using a finite impulse response filter. Noisy channels were identified and removed using automated procedures. EEG data was subsequently segmented into two second epochs, and epochs containing noisy data were identified as follows: the mean activity of all EEG channels was computed, and the threshold was set at ± 2 times the standard deviation of the mean activity. Epochs containing activity exceeding the threshold value were marked and subsequently removed. This left a total of 160 (± 15) two second epochs in the stroke cohort and a total of 73 (± 12) in the MS cohort. To identify and remove ocular movements and blink artifacts from the EEG data, an independent component analysis (ICA) implemented within EEGLAB was used. ICA is a blind source decomposition algorithm that enables the separation of statistically independent sources from multichannel data^37^. The components were subsequently visually inspected and those containing ocular movements or blink artifacts were removed. The previously removed channels were then interpolated back into the dataset and finally, the EEG data was re-referenced against the grand average of all scalp electrodes.

### Graph Theory Estimates

#### Functional Connectivity Analysis

EEG connectivity analysis was carried out using the exact low-resolution electromagnetic tomography (eLORETA) software (The KEY Institute of Brain-Mind Research University Hospital of Psychiatry, Zurich; http://www.uzh.ch/keyinst/NewLORETA/LORETA01.htm). The eLORETA algorithm is a well-established linear inverse solution for EEG signals^38^.

Following whole brain sources reconstruction, connectivity was computed using the eLORETA software on four brain regions, divided into motor and sensory networks of the left and right hemisphere based on Broadmann areas (BAs). Each BA is a region of interest (ROI). The BAs that formed the motor network for both the left and right hemisphere included BA4, 6, 8, 9, 24 and 32, while the BAs that formed the sensory network for both the left and right hemisphere included BA1, 2, 3, 5, 7, 40 and 43.

Current density time series of all BAs within each of the four networks was computed in eLORETA and used to estimate the functional connectivity using the Lagged Linear Coherence (LagR) algorithm, not affected by volume conductance and low spatial resolution in each of the four networks^39^. As the brain functional coherence/connectivity is frequency-dependent, the Lagged Linear Coherence was computed for each of the 5 independent frequency bands of delta (2-4 Hz), theta (4-8 Hz), alpha (8-13 Hz), beta (13-30 Hz) and gamma (30-45 Hz).

#### Graph Analysis

A network is a mathematical representation of a real-world complex system and is defined by a collection of nodes (vertices) and links (edges) between pairs of nodes. Nodes in large-scale brain networks represent brain regions, while links represent anatomical or functional connections. Nodes should ideally represent brain regions with coherent patterns of anatomical or functional connections. The connectivity parameters extracted between all pairs of ROIs for each frequency band is in the form of a square matrix W, with dimensions equal to the number of ROIs. Each row and column within matrix W represent nodes, while the values within the matrix represent the strength of connection between each pair of nodes.

Once the networks of interest were constructed, the core measures of graph theory that summarize the aspects of segregation and integration of a network were computed using the Brain Connectivity Toolbox^29^. Segregation refers to the degree to which network elements form individual and separate clusters and is measured by the clustering coefficient (*C*). Integration refers to the capacity of the network to become interconnected and exchange information and is measured by the parameter characteristic path length (*L*). The clustering coefficient and characteristic path length represent the efficiency of the network with respect to local and global connectedness respectively. Weighted clustering (Cw) coefficient and weighted characteristic path length (Lw) were computed as a measure of segregation and integration of the network as follows:

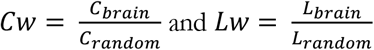

Where C_brain_ and L_brain_ are the clustering coefficient and characteristic path length derived from the connectivity matrix of each participant. C_random_ and L_random_ are the mean values of the clustering coefficient and characteristic path length of 100 surrogate random networks that have the same basic characteristics as the original network. A measure of network small-worldness (Sw) was therefore defined as the ratio between Cw and Lw; the ratio between local connectedness and the global integration of the network.

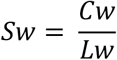

When Sw has a value of approximately 1, a network is said to have “small-world properties” meaning a good combination of high levels of local clustering among nodes and proper paths that globally link all network nodes (all nodes of a large system are linked through relatively few intermediate steps). Sw values greater than 1 suggest high levels of local clustering among nodes and many short paths that globally link all nodes of the network, while Sw values less than 1 suggest poor local connectivity and stunted connections.

### Statistical Analysis

All statistical analysis was performed using R (RStudio Version 1.2.5033). Spearman rank correlations were used to identify the association between trait fatigue (FSS-7 or mFIS scores) and all continuous demographic measures (age, grip strength, NHPT, HADS – Depression, HADS – Anxiety, Time Post-Stroke, EDSS and BDI). Wilcoxon rank sum tests were used to identify the association between trait fatigue (FSS-7 or mFIS scores) and all categorical demographic measures (sex, hemisphere affected, type of stroke and vascular territory affected, MS type). The distribution of the dependent variable was assessed using the Shapiro-Wilk’s test of normality, while the homogeneity of variances was assessed using the Levene’s test.

In the case of normally distributed variables, a two-way mixed ANOVA was performed to evaluate the effects of the between-subject factor Fatigue group (Stroke_Low_, Stroke_High_, MS_High_) and the within-subject factors Frequency band (delta, theta, alpha, beta, gamma) on small-worldness (dependent variable), for each of the two networks (sensory and motor) and each hemisphere (left and right). Greenhouse-Geisser epsilon adjustment was employed in these analyses to correct for any deviations from sphericity. Post-hoc tests were subsequently used to simple main effects and pairwise comparisons (pairwise t-tests) with Bonferroni adjustment.

## Results

### Participant Demographics

Twenty-nine stroke survivors completed the study (11 females and 18 males). The median FSS-7 score was 5.29 (IQR = 2.57) in females and 2.50 (IQR = 2.46) in males. The Wilcoxon test showed that the difference in FSS-7 score was marginally non-significant (p = 0.05, effect size = 0.37). Spearman rank correlations between trait fatigue (FSS-7) and all continuous demographic measures revealed a significant positive association between trait fatigue and HADS-Depression (Spearman ρ = 0.41, p = 0.03), while no other variable correlated with trait fatigue.

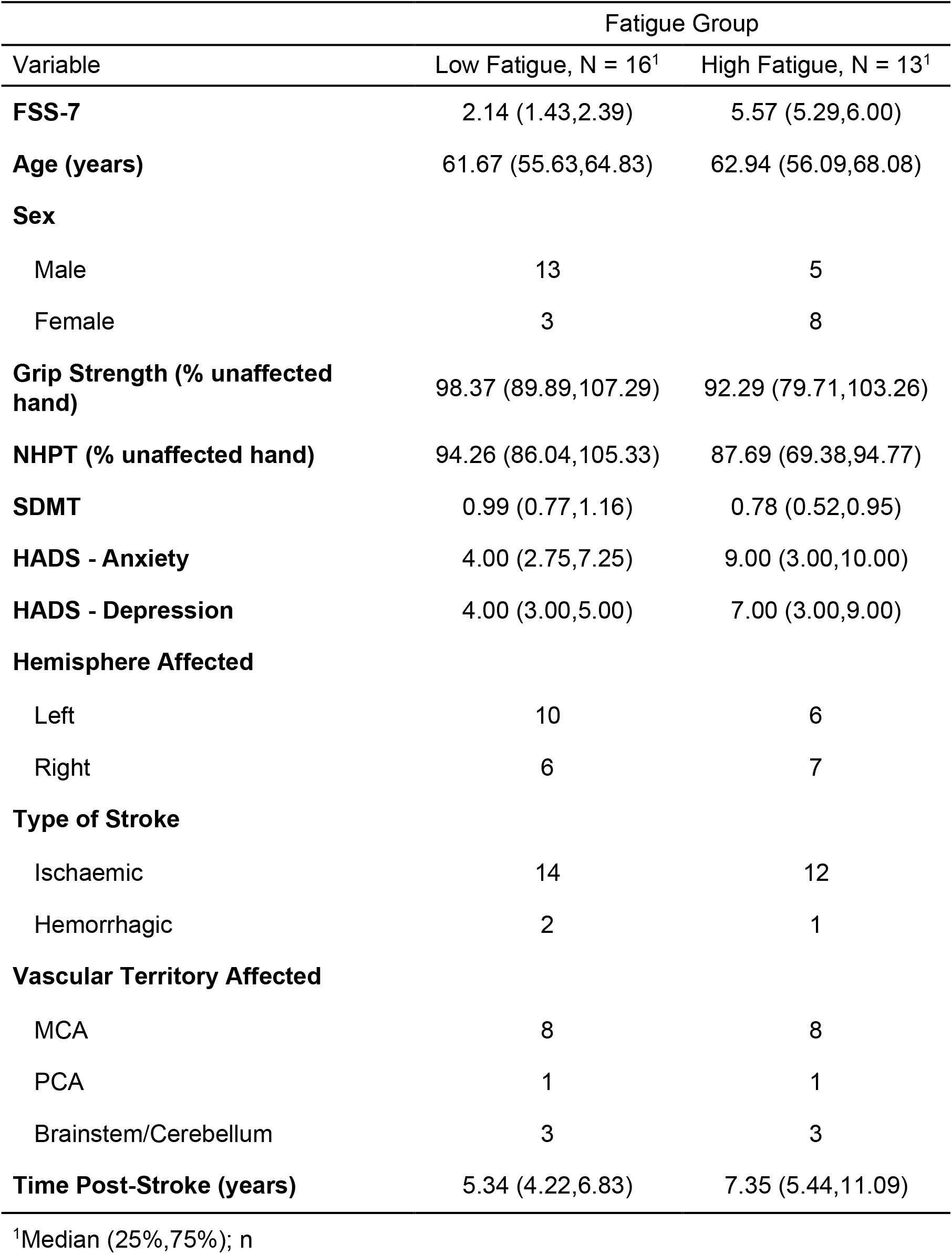

| Variable | Fatigue Group |  |
| --- | --- | --- |
|  | Low Fatigue, N = 16<sup>1</sup> | High Fatigue, N = 13<sup>1</sup> |
| **FSS-7** | 2.14 (1.43,2.39) | 5.57 (5.29,6.00) |
| **Age (years)** | 61.67 (55.63,64.83) | 62.94 (56.09,68.08) |
| **Sex** |  |  |
| Male | 13 | 5 |
| Female | 3 | 8 |
| **Grip Strength (% unaffected hand)** | 98.37 (89.89,107.29) | 92.29 (79.71,103.26) |
| **NHPT (% unaffected hand)** | 94.26 (86.04,105.33) | 87.69 (69.38,94.77) |
| **SDMT** | 0.99 (0.77,1.16) | 0.78 (0.52,0.95) |
| **HADS - Anxiety** | 4.00 (2.75,7.25) | 9.00 (3.00,10.00) |
| **HADS - Depression** | 4.00 (3.00,5.00) | 7.00 (3.00,9.00) |
| **Hemisphere Affected** |  |  |
| Left | 10 | 6 |
| Right | 6 | 7 |
| **Type of Stroke** |  |  |
| Ischaemic | 14 | 12 |
| Hemorrhagic | 2 | 1 |
| **Vascular Territory Affected** |  |  |
| MCA | 8 | 8 |
| PCA | 1 | 1 |
| Brainstem/Cerebellum | 3 | 3 |
| **Time Post-Stroke (years)** | 5.34 (4.22,6.83) | 7.35 (5.44,11.09) |

<sup>1</sup>Median (25%,75%); n

Seventeen MS patients completed the study (13 females and 4 males). The median mFIS score was 51 (IQR = 16) in females and 39.5 (IQR = 4.5) in males. The Wilcoxon test showed that the difference in mFIS score was non-significant (p = 0.11, effect size = 0.40). Spearman rank correlations between trait fatigue (mFIS) and all continuous demographic measures (age, EDSS and BDI) revealed no significant associations.

| Variable | N = 17<sup>1</sup> |
| --- | --- |
| **mFIS** | 50.00 (39.00,57.00) |
| **Age (years)** | 41.00 (33.00,46.00) |
| **Sex** |  |
| Male | 4 |
| Female | 13 |
| **Expanded Disability Status Scale** | 1.00 (0.00,1.50) |
| **Beck Depression Inventory** | 10.00 (8.00,12.00) |
| **MS Type** |  |
| Relapsing Remitting | 16 |
| Primary Progressive | 1 |
| **Disease Duration** | 6.00 (1.10,11.10) |
| **Annual Relapse Rate** |  |
| 0 | 12 |
| 1 | 4 |
| 2 | 1 |
| **Disease Modifying Therapies** |  |
| Azatioprina | 1 |
| Fingolimod | 2 |
| INF beta1a | 3 |
| INF beta1aHD | 1 |
| INF beta1b | 6 |
| Na | 2 |
| Natalizumab | 1 |

<sup>1</sup>Median (25%,75%); n

### Clinical Characteristics of Stroke and MS

The association between trait fatigue (FSS-7) and the clinical characteristics of the stroke was assessed across all stroke survivors. The median FSS-7 score in those with right hemisphere strokes was 4.43 (IQR = 3.00) and 2.71 (IQR = 3.75) in those with left hemisphere strokes (Wilcoxon test: p = 0.50, effect size r = 0.13). The median FSS-7 score in those with ischemic strokes was 2.93 (IQR = 3.46) and 3.86 (IQR = 2.29) in those with hemorrhagic strokes (Wilcoxon test: p = 0.51, effect size r = 0.13). Regarding the vascular territory affected, the data from five stroke survivors was missing as the clinical notes could not be retrieved. The median FSS-7 score in those where the MCA was affected was 3.36 (IQR = 3.25), the median FSS-7 score in those where the PCA was affected was 3.07 (IQR = 2.07), while the median FSS-7 score in those where the Brainstem/Cerebellum was affected was 4.21 (IQR = 2.68) (Kruskal-Wallis test: p = 0.57, effect size η^2^ = −0.04). A spearman rank correlation between FSS-7 and the Time Post-Stroke at which the participants took part in the study showed no significant association (spearman ρ = 0.08, p = 0.67). Any meaningful interpretation of the effect of the type of stroke and vascular territory affected on FSS-7 in the current cohort of stroke survivors is difficult given the skewed numbers.

The association between trait fatigue (mFIS) and the clinical characteristics of MS was assessed across all MS patients. The median mFIS score in those with relapsing remitting MS was 50 (IQR = 18.2), whereas the mFIS score in the one patient with primary progressive MS was 40 (Wilcoxon test: p = 0.61, effect size r = 0.11). A spearman rank correlation between mFIS and the disease duration showed no significant association (spearman ρ = −0.37, p = 0.14). The median mFIS score in those with an ARR of zero was 43.5 (IQR = 17.2), the median mFIS score in those with an ARR of one was 54 (IQR = 8.3), while the mFIS score in the one patient with an ARR of two was 38 (Kruskal-Wallis test: p = 0.18, effect size = 0.10).

### Small Worldness of Sensory Network

Two separate two-way mixed ANOVA, one for each hemisphere, with the within-subject factor of Frequency band (delta, theta, alpha, beta, gamma) and the between-subject factor of Fatigue group (Stroke_Low_, Stroke_High_, MS_High_) was performed. Across the five frequency bands and two hemispheres in the three fatigue groups (460 data points) there were twelve extreme outliers (6 delta, 1 theta and 1 gamma in the right hemisphere; 1 delta, 2 theta and 1 beta in the left hemisphere). Following removal of outliers, the data was normally distributed, as assessed by the Shapiro-Wilk’s test of normality (p > 0.05) and there was homogeneity of variances (p > 0.05) as assessed by Levene’s test of homogeneity of variances. The ANOVA in the right hemisphere revealed a main effect of Frequency band (F_(2.95,103)_ = 3.38, p = 0.02, η^2^ = 0.07), while there was no effect of fatigue and no two-way interaction between fatigue and frequency band. Post-hoc multiple pairwise comparisons revealed a significant difference in beta band frequency power (Figure 1B) between the Stroke_Low_ and Stroke_High_ groups, as well as between Stroke_Low_ and MS_High_ groups in the beta frequency band (p = 0.02 and p = 0.04 respectively). In the left hemisphere, there was no effect of fatigue, frequency band or interaction between fatigue and frequency band.

**Figure 1.**
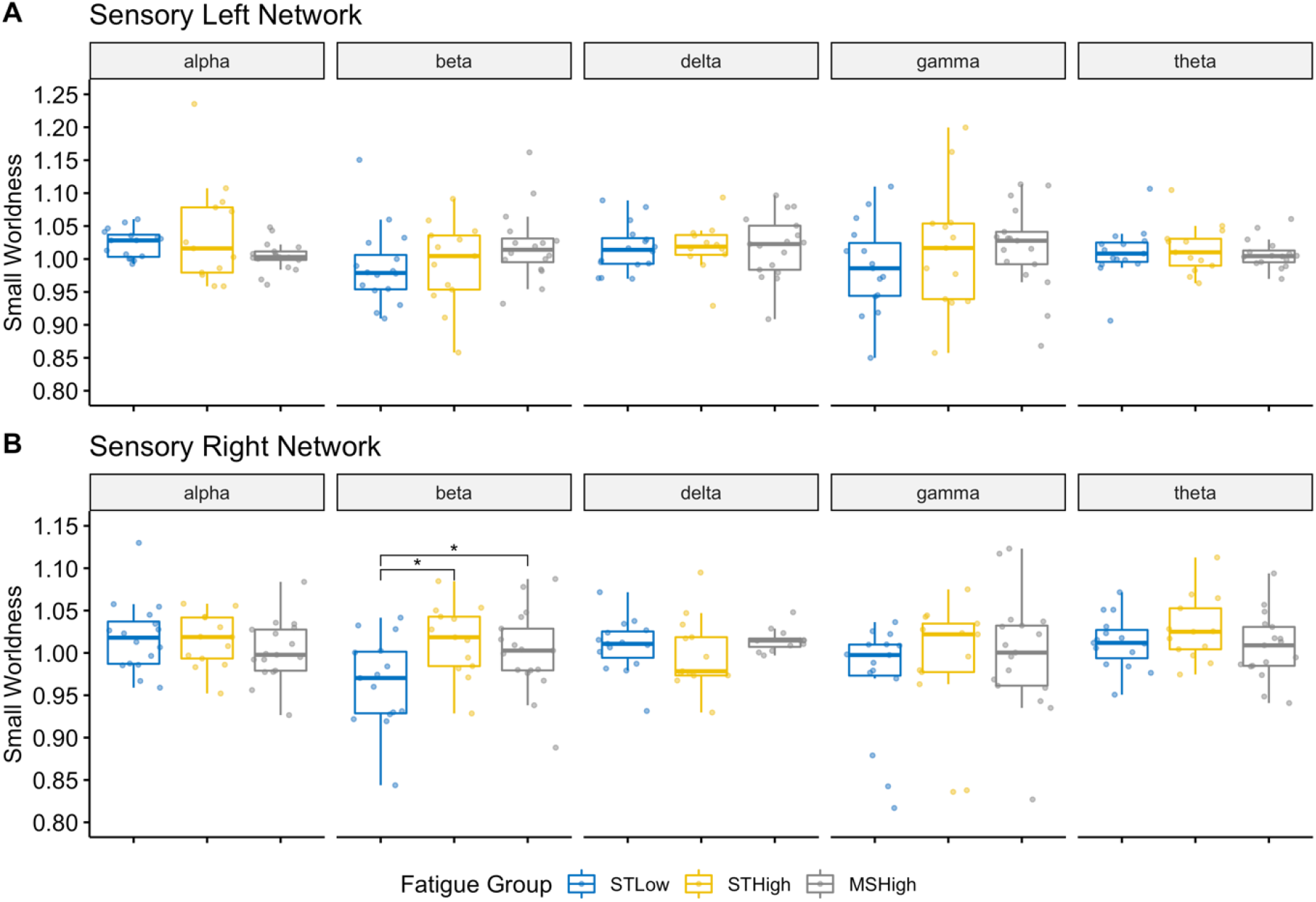
Functional Connectivity of the Sensory Network. The value Small Worldness across the three fatigue groups (Stroke_Low_ in blue, Stroke_High_ in yellow, MS_High_ in grey) is displayed using boxplots across the five frequency bands (alpha, beta, delta, gamma and theta) for the sensory network in the left (A) and right (B) hemisphere. Significant differences between fatigue groups are indicated using asterisks (* = p < 0.05)

### Small Worldness of Motor Network

Two separate two-way mixed ANOVA, one for each hemisphere, with the within-subject factor of Frequency band (delta, theta, alpha, beta, gamma) and the between-subject factor of Fatigue group (Stroke_Low_, Stroke_High_, MS_High_) was performed (Figure 2). Across the five frequency bands and two hemispheres in the three fatigue groups (460 data points) there were ten extreme outliers (1 delta, 2 beta and 2 gamma in the right hemisphere; 1 delta and 4 gamma in the left hemisphere). The data was normally distributed, as assessed by the Shapiro-Wilk’s test of normality (p > 0.05) and there was homogeneity of variances (p > 0.05) as assessed by Levene’s test of homogeneity of variances. The ANOVA of the left hemisphere revealed a main effect of fatigue (F_(2,38)_ = 8.49, p < 0.001, η^2^ = 0.07), while there was no effect of frequency band and no interaction between fatigue and frequency band. In the right hemisphere, there was no effect of fatigue, frequency band or interaction between fatigue and frequency band. The main effect of fatigue in the left hemisphere was followed up by multiple pairwise comparisons to determine in which frequency band the group means differed. The mean Sw score was significantly different between the Stroke_Low_ and Stroke_High_ groups in the beta and theta frequency bands of the left hemisphere (p = 0.03 and p = 0.03 respectively; Figure 2A).

**Figure 2.**
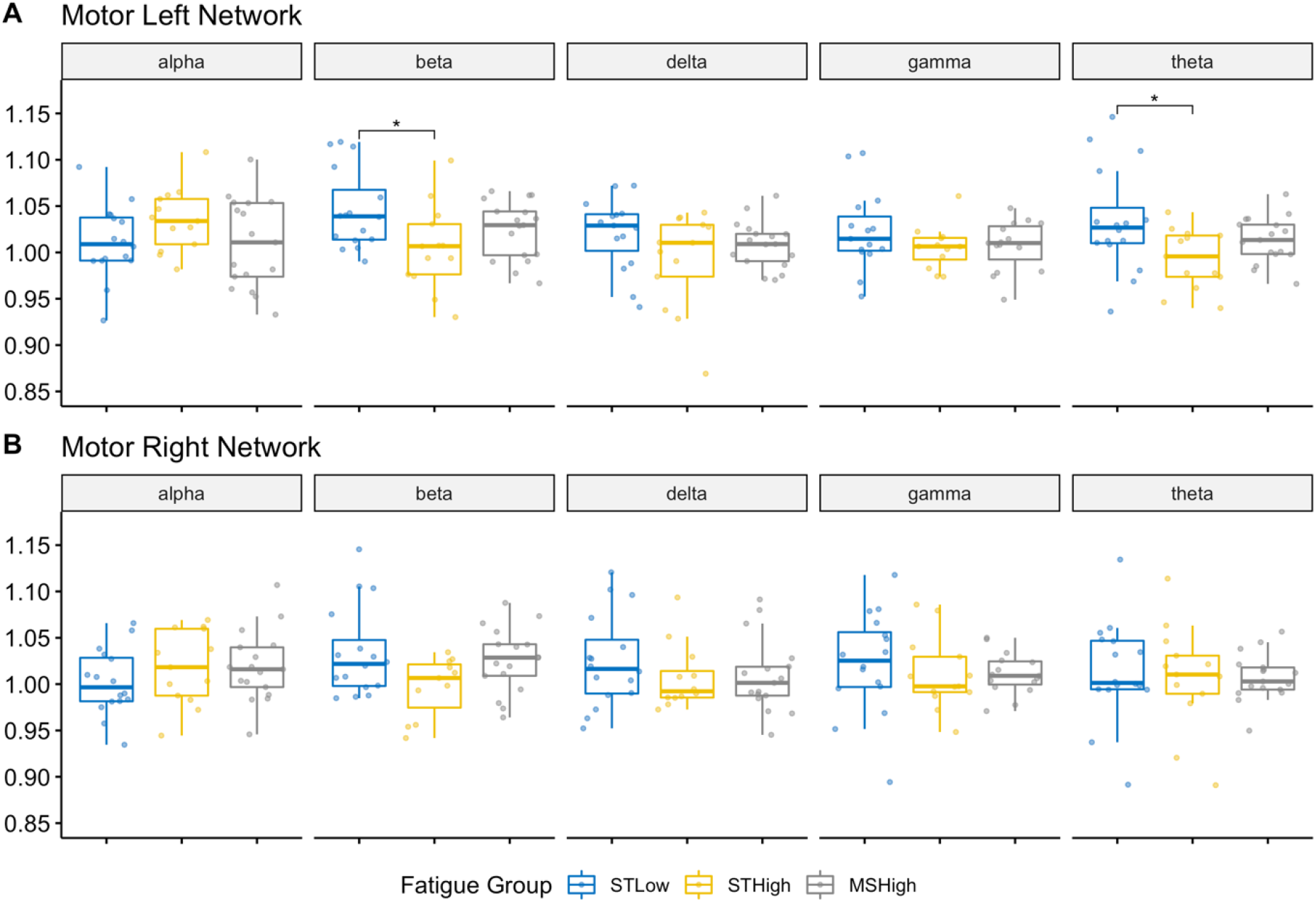
Functional Connectivity of the Motor Network. The value Small Worldness across the three fatigue groups (Stroke_Low_ in blue, Stroke_High_ in yellow, MS_High_ in grey) is displayed using boxplots across the five frequency bands (alpha, beta, delta, gamma and theta) for the sensory network in the left (A) and right (B) hemisphere. Significant differences between fatigue groups are indicated using asterisks (* = p < 0.05)

**Figure 3:**
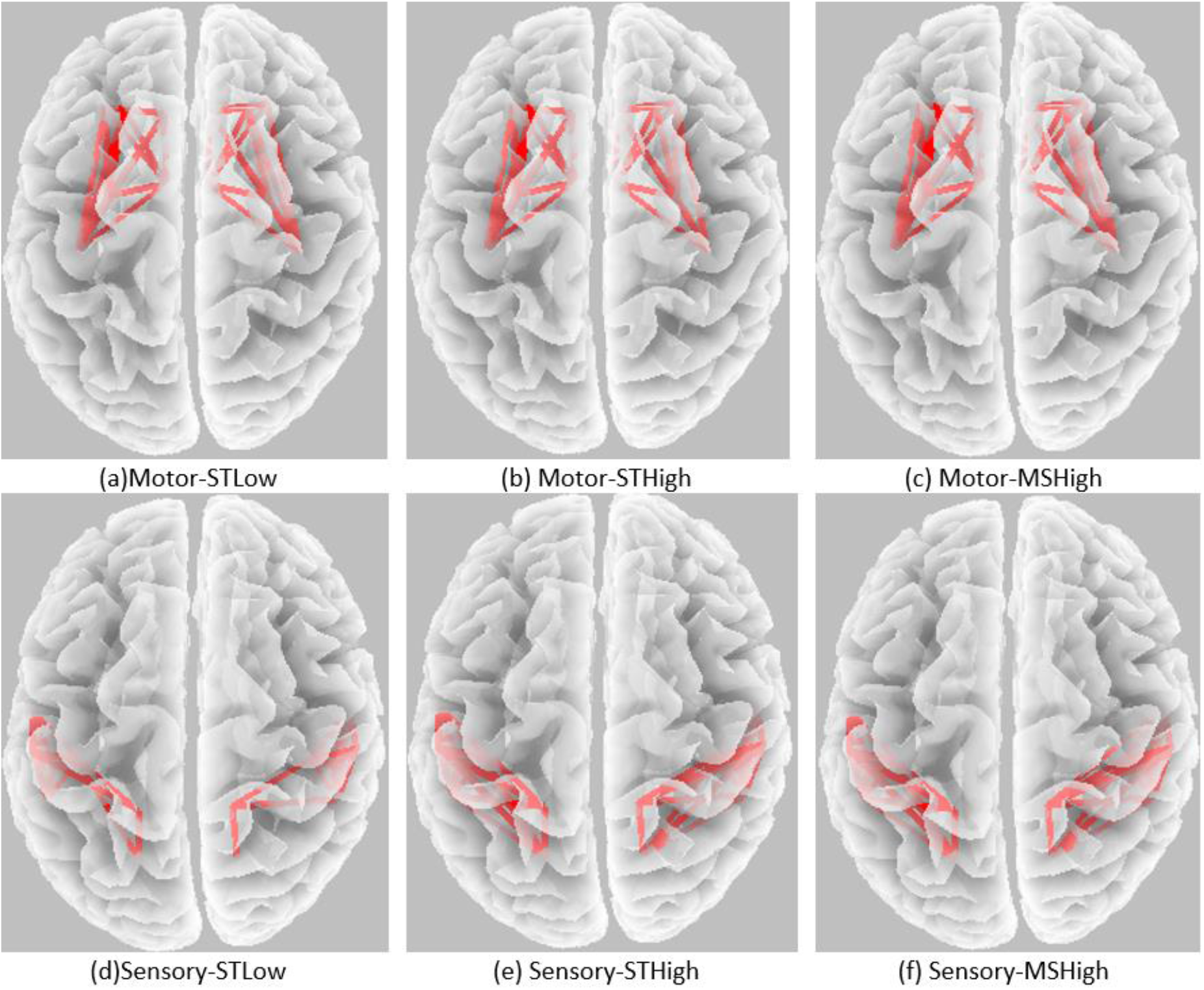
A visual representation of the small worldness index in the motor (a,b,c) and sensory (d,e,f) in the right and left hemispheres of MS (a,d) and stroke (b,c,e,f) groups.

## Discussion

In forty-six patients with either stroke or MS, we show that those with high levels of trait fatigue exhibited significantly higher levels of small worldness in the right sensory network in the beta band frequency. Additionally, only in the stroke survivor group, those with high fatigue showed significantly lower levels of small worldness in the left motor network, both in the beta and theta frequency bands. There was no association between clinical features of stroke or MS and trait fatigue. Such a lack of association has been previously reported in both stroke^1^ and MS^40^. As investigated by small-worldness, functional brain connectivity simultaneously reconciles the opposing demands of functional integration and segregation. The small-world index reflects the balance of functionally specialized (segregated) modules with a robust number of intermodular (integrating) links. Here we observed that fatigue is paralleled by the alteration of small-world organization in sensory areas.

Beta band activity is commonly known as the sensorimotor ‘idling’ rhythm seen in all cortical and sub-cortical motor areas at rest. Movement desynchronises beta band oscillations which lead to the idea that beta frequency is the rhythm of rest for motor areas. However, recent proposals suggest that beta band activity may not simply reflect a lack of movement but is rather an indicator for maintenance of sensorimotor status quo^41^. During periods of spontaneous enhancement in resting beta band activity, movements are slower, than when resting beta activity is lower^42^. Such baseline beta activity influence on movement speed indicates that resting beta band activity may signal maintenance of sensorimotor status quo. In another study, entraining motor cortex at 20 Hz frequency resulted in slower movements without any effect on reaction times^43^. In movement disorders such as Parkinson’s disease that is defined by slow movements, an abnormally high resting state beta rhythm is seen in sub-cortical motor structures^44^, with stimulation aimed at desynchronising resting beta rhythms temporarily, speeding up movements. In post-stroke fatigue, while there is no difference in reaction times, there is slowing of movements^19^, perhaps because there is a resistance to change sensorimotor status quo as reflected by enhanced beta rhythm small-worldness shown in this study.

Beta band mediates the synchronization with muscular activity as indexed by the cortico-muscular coherence (CMC), whose extensive reliability has been recently reviewed^45^, indicating beta band CMC as reflecting the interaction and the flow of information between the cerebral cortex sending commands to muscle tissue and the afferent feedback from the controlled districts. Sensory and proprioceptive feedback is inherently part of motor control^46,47^ and modulates the CMC^48,49^, with about 30% of the corticospinal tract fibers originating from the primary somatosensory areas^50^. In terms of resting state oscillatory activity, it has been observed that primary somatosensory area prevails in alpha and low beta, while primary motor area in gamma, both from non-invasive recordings^51^ and stereo-EEG investigation^52^. It is well-established that those with fatigue, both in stroke and MS, have little relation to motor or cognitive impairment, thereby such limitations cannot explain poor behavioural flexibility. The question then arises, what drives poor behavioural flexibility. To answer this, we must look to what is represented by beta band activity. From a predictive coding perspective of brain processing, oscillatory brain activity in the beta band represents sensory ‘priors’ related to expected movement related sensory states of the body^53^. Movement related sensory priors in disease conditions with high fatigue, is informed by movements that are perceived as highly effortful^21^. At rest, such high effort movement priors give rise to the feeling of fatigue^3^. Therefore, the driver for increased resting state beta band activity is directly linked to what is represented by beta band oscillations of the predictive brain. To avoid experiencing a state of high effort in the future, the fatigue brain seeks to maintain its sensorimotor status quo by increasing its resting state beta band activity.

In this study we provide first evidence for a common mechanism of pathological fatigue in two neurological conditions that present with fatigue as a significant symptom, stroke and MS. Both in post-stroke and MS fatigue, there are several reports of altered resting state connectivity^14,54–58^. In stroke, suggestions of parietal hypoconnectivity and frontal hyper connectivity^58^ with reversed inter-hemispheric balance of connectivity^14^, are implicated in manifestation of fatigue. In MS, changes in default mode network^56^ and involvement of striatal circuits involved in movement, sensation and motivation^55^ have all been implicated in development of fatigue. While several brain regions have been implicated in both diseases as the core regions involved in fatigue, very few of the studies performed hypothesis driven analysis on resting state activity. In a previous study on a different fatigued MS cohort, a functional connectivity alteration at rest occurred in the sensory cortical networks, mediated by beta-band oscillatory activity^25^ in good agreement with present results. In the same line of observations, in MS an intracortical connectivity index within the hand somatosensory representation, coding for sensorimotor dexterity^59^, appeared distorted^60^.

In the present study we hypothesised that processing of sensory information was at the heart of pathological fatigue irrespective of disease processes. Investigating sensory and motor networks, we revealed that both in MS and Stroke, a common mechanism of altered connectivity in the sensory networks underlies symptoms of fatigue. This disease independent mechanism strengthens the idea that chronic fatigue may be triggered, but not maintained by the disease condition, with chronic fatigue resulting from maladaptive sensorimotor feedback functionality and plastic changes.

Limitations of this study include the lack of low fatigue MS group, and the relatively small number of participants in both stroke and MS groups. This investigation should be viewed as a proof of concept study demonstrating a common mechanism across disease states, rather than a study revealing new mechanisms that underpin fatigue. This study paves the way for future studies to treat pathological fatigue in chronic neurological conditions as driven by common factors, and therefore design large comparative studies of fatigue across diseases.

In summary, we show that in two neurological diseases with different aetiologies, similar pathological connectivity in sensory networks explains high levels of self-reported fatigue. The frequency band in which the alteration manifested, beta frequency band, suggests that high effort sensory representations of movement might underlie pathological fatigue.

## Notes

### Competing Interest Statement

The authors have declared no competing interest.

